# Insights into kinetics, release, and behavioral effects of brain-targeted hybrid nanoparticles for cholesterol delivery in Huntington’s disease

**DOI:** 10.1101/2020.11.24.395525

**Authors:** Giulia Birolini, Marta Valenza, Ilaria Ottonelli, Alice Passoni, Monica Favagrossa, Jason T. Duskey, Mauro Bombaci, Maria Angela Vandelli, Laura Colombo, Renzo Bagnati, Claudio Caccia, Valerio Leoni, Franco Taroni, Flavio Forni, Barbara Ruozi, Mario Salmona, Giovanni Tosi, Elena Cattaneo

**Author notes:** co-first authors (Giulia Birolini, Marta Valenza).

## Abstract

Supplementing brain cholesterol is emerging as a potential treatment for Huntington’s disease (HD), a genetic neurodegenerative disorder characterized, among other abnormalities, by inefficient brain cholesterol biosynthesis. However, delivering cholesterol to the brain is challenging due to the bloodbrain barrier (BBB), which prevents it from reaching the striatum, especially, with therapeutically relevant doses.

Here we describe the distribution, kinetics, release, and safety of novel hybrid polymeric nanoparticles made of PLGA and cholesterol which were modified with an heptapeptide (g7) for BBB transit (hybrid-g7-NPs-chol). We show that these NPs rapidly reach the brain and target neural cells. Moreover, deuterium-labeled cholesterol from hybrid-g7-NPs-chol is released in a controlled manner within the brain and accumulates over time, while being rapidly removed from peripheral tissues and plasma. We confirm that systemic and repeated injections of the new hybrid-g7-NPs-chol enhanced endogenous cholesterol biosynthesis, prevented cognitive decline, and ameliorated motor defects in HD animals, without any inflammatory reaction.

In summary, this study provides insights about the benefits and safety of cholesterol delivery through advanced brain-permeable nanoparticles for HD treatment.

## 1. Introduction

Targeting the brain with therapeutic agents is made difficult by the blood-brain barrier (BBB). Current strategies to circumvent this problem include temporary BBB disruption, conjugation of brain-permeable ligands to the drug of interest, intranasal delivery, or direct delivery of molecules into the brain by means of invasive strategies [1]. Among them, nanocarriers [2] decorated on their surface with brain-targeting ligands [3–4] are emerging because of their non-invasiveness. Polymeric nanoparticles (NPs) made of the FDA-approved polymer poly (D,L-lactide-co-glycolide) (PLGA) and surface-engineered with the g7 glycopeptide (Gly-L-Phe-D-Thr-Gly-L-Phe-L-Leu-L-Ser(O-beta-D-glucose)-CONH2) are reported to transport molecules into the CNS after their systemic administration in rodents [5]. g7 stimulates membrane curvature and, following endocytosis of the whole carrier at the BBB, it promotes BBB transit by multiple pathways [6]. Previous studies indicate that about 10% of the injected g7-NP reaches the brain [6–11].

This system has been applied to deliver drugs to cope with Huntington’s disease (HD), a neurodegenerative genetic disorder caused by a CAG repeat expansion in the gene encoding the Huntingtin protein and characterized by motor, cognitive, and psychiatric disturbances [12]. In particular, in a first study, systemic injections of PLGA-g7-NPs loaded with cholesterol (PLGA-g7-NPs-chol) were found to prevent synaptic and cognitive defects in a transgenic mouse model of HD [13]. The rationale behind this treatment stands on the solid demonstration of a marked reduction in cholesterol biosynthesis in the rodent HD brain [14–18]. In a subsequent study, cholesterol was infused for 4 weeks into the striatum of HD mice through osmotic mini-pumps, a well-characterized system for controlled drug delivery. This led to the identification of the optimal dose of cholesterol needed to restore synaptic and neuropathological features and reverse motor and cognitive abnormalities in HD mice [19]. Recently, the effectiveness of nose-to-brain delivery of cholesterol to the HD brain has also been tested [20]. In the first study with PLGA-g7-NPs-chol, the amount of cholesterol delivered to the brain was sub-optimal compared with the dose identified with osmotic mini-pumps. In particular, this kind of NPs was estimated to deliver to the brain about 2 μg of cholesterol in each systemic injection [13], while about 13 μg of cholesterol per day were infused in the striatum with mini-pumps, enabling the maximum benefit in HD mice following 4 weeks of treatment [19].

The development of a more efficient, non-invasive, brain-targeted cholesterol-based therapy for HD patients requires an optimized generation of NPs with improved pharmaceutical properties. In particular, in this study we tested the therapeutic value of new nanoparticles whose formulation process, structural characteristics, and drug loading capacity have been enhanced.

These NPs, herein named *hybrid-g7-NPs-chol,* were prepared using a novel formulation that efficiently combines materials (g7-PLGA, PLGA, and cholesterol as starting materials for formulation) [21], using nanoprecipitation (MIX-N) or single emulsion (MIX-SE) with surfactants such as Pluronic F68 in MIX-N and polyvinyl alcohol (PVA) in MIX-SE. The maximal percentages of cholesterol present in hybrid-g7-NPs-chol MIX-N and MIX-SE were 32.7 ± 2% and 41.5 ± 1.5%, respectively, reaching a cholesterol content which is around 40 times higher than that of PLGA-g7-NPs-chol [13]. This drug delivery system is therefore expected to deliver to the HD brain a therapeutic dose of cholesterol closer to that identified with the osmotic mini-pumps [19], while being less invasive and more translatable. This study therefore aimed to investigate the distribution of hybrid-g7-NPs-chol in different brain regions, peripheral tissues, and the circulation, and the kinetics of cholesterol release, the benefits to animals’ behavior. Potential side effects which might emerge following chronic treatment in HD mice were also investigated.

## 2. Method Section

### 2.1 Production and characterization of hybrid-g7-NPs-chol

Starting from already published results [13, 21], we produced hybrid NPs using a nanoprecipitation method (N) and simple emulsion (SE). Following previous readouts on the polymer ratio to be used in formulation [21], 1:1 w/w ration between PLGA and Cholesterol was adopted. In details, all hybrid nanoparticles were formulated starting from 50 mg of a mixture of Cholesterol (Sigma-Aldrich, Milan, Italy), PLGA (PLGA R503H Evonik, Essen, Germany) and PLGA-g7 in a weight ratio of 1:0.8:0.2 (25mg Chol, 20mg PLGA, 5mg PLGA-g7).

PLGA-g7 was synthesized via amide formation in the Nanotech Laboratory of the University of Modena and Reggio Emilia, following a previously described protocol [8, 26]. Pure g7 was purchased from Mimotopes (Clayton, Victoria, Australia).

To obtain **MIX-N**, Chol and PLGA mixture was dissolved in acetone (4 mL). The organic phase was then added dropwise into 50 mL of a 0.5% (w/v) Pluronic-F68 aqueous solution at 45 °C under magnetic stirring (1300 rpm). After 20 min, the organic solvent was removed at 30 °C under reduced pressure (10 mmHg). The MIX-NPs were recovered and purified three times by an ultracentrifugation process carried out at 14,500 rpm for 10 min (4 °C; Sorvall RC28S, Dupont, Brussels, Belgium) to remove the unformed material and the free surfactant fraction in the solution. The obtained MIX-NPs were re-suspended in 3 mL of a 0.2% (w/v) Pluronic-F68 aqueous solution at room temperature and gently sonicated until completely resuspended.

As cryoprotectant, 150 mg of threalose (Sigma-Aldrich, Milan, Italy) dissolved in 0,5mL of 0.2% Pluronic-F68 solution were added to the NPs suspension before flash freezing with dry ice and methanol bath. NPs were stored at −20°C until use.

To obtain **MIX-SE**, Chol:PLGA mixture were dissolved in dichloromethane (4mL) and emulsified in 20 mL of 1% (w/v) PVA aqueous solution by sonication (Microson Ultrasonic cell disruptor, Misonix Inc. Farmingdale, NY, USA) (80 W over 1 min) under cooling (5 °C). Then, the O/W emulsion was stirred for at least 3 h (1300 rpm; RW20DZM, IKALabortechnik, Staufen, Germany) at r.t. to allow the solvent evaporation. The MIX-SE were collected and purified by ultracentrifugation as previously described for MIX-N, and stored at 4 °C before the use.

Fluorescently labelled MIX NPs were produced with the same protocol described in 1.1.1, adding 2% in weight of Cy5 derived PLGA and 2% in weight of Bodipy-Cholesterol (Avanti, Alabama, USA). Cy5-PLGA was synthesized in the Nanotech Laboratory of the University of Modena and Reggio Emilia using a protocol published in [27]. The total composition was a mixture of Chol:Chol-Bodipy:PLGA:PLGA-Cy5:PLGA-g7 in weight ratio of 0.96:0.04:0.76:0.04:0.2 (24 mg Chol, 1 mg Chol-Bodipy, 19 mg PLGA, 1 mg PLGA-Cy5, 5 mg PLGA-g7). After centrifugation, NPs were resuspended in an aqueous solution of Pluronic-F68 2% and added of threalose as previously reported before flash freezing.

g7-NPs-d6-Chol were obtained using the same protocol described in paragraph 1.1.1. For the formulation, 50 mg of a mixture of D6 Chol:PLGA:PLGA-g7 in weight ratio of 1:0.8:0.2 (25 mg D6 Chol, 20 mg PLGA, 5 mg PLGA-g7) were used.

### 2.2 Production of PLGA-g7-NPs loaded with Cholesterol

PLGA-g7-NPs-chol were produced as reported in [13], confirming technological and pharmaceutical features as described in Table S1 as shown before.

### 2.3 Chemico-physical and morphological characterization

100μl of each type of NPs suspension was freeze-dried (−60 °C, 1 · 10-3 mm/Hg for 48 h; LyoLab 3000, Heto-Holten, Allerod, Denmark) and the yield (Yield%) was calculate as follows: Yield (%)=[(mg of freeze dried MIX-N or MIX-SE)/(mgPLGA+mg Chol)]×100 Mean particle size (Z-Average) and polydispersity index (PDI) of the samples were determined using a Zetasizer Nano ZS (Malvern, UK; Laser 4 mW He-Ne, 633 nm, Laser attenuator Automatic, transmission 100-0.0003%, Detector Avalanche photodiode, Q.E.>50% at 633 nm). Samples were diluted in MilliQ water at about 0.1 mg/mL. The results were also expressed as intensity distribution, i.e. the size 10% [D(10)], 50% [D(50)] and 90% [D(90)], below which all the MIX NPs are placed. The zeta potential (ζ-pot l) was measured using the same equipment with a combination of laser Doppler velocimetry and phase analysis light scattering (PALS). All data are expressed as means of at least three determinations carried out for each prepared lot (three lots for each sample).

The morphology of the samples was evaluated by atomic force microscopy (AFM) (Park Instruments, Sunnyvale, CA, USA) and scanning transmission electron microscopy (STEM) as reported in [21], confirming the same results as reported in supplementary figure S1.

AFM analysis were conducted at about 20 °C operating in air and in non-contact mode using a commercial silicon tip-cantilever (high resolution noncontact “GOLDEN” Silicon Cantilevers NSG-11, NT-MDT, tip radius 10 nm; Zelenograd, Moscow, Russia) with stiffness of about 40 Nm-1 and a resonance frequency around 160 kHz. A drop of each hybrid NPs suspension was diluted with distilled water (about 1:5 v/v) before application on a small mica disk (1 cm×1 cm).

After 2 minutes, the excess of distilled water was removed using a paper filter and the sample analyzed. Two kinds of images were obtained: the first is a topographical image and the second is indicated as “error signal”. This error signal is obtained by comparing two signals: the first one representing the amplitude of the vibrations of the cantilever, and the second the amplitude of a reference point. The images obtained by this method show small superficial variations of the samples. Images were processed and analyzed using software from Gwyddion (Department of Nanometrology, Czech Metrology Institute, Brno, Czech Republic).

The internal structure/architecture of the samples was analyzed by scanning transmission electron microscopy (STEM). Briefly, a drop of a water-diluted suspension of the samples (about 0.03 mg/mL) was placed on a 200-mesh copper grid (TABB Laboratories Equipment, Berks, UK), allowed to adsorb, and the suspension surplus was removed by filter paper. All grids were analyzed using a Nova NanoSEM 450 (FEI, Oregon, USA) transmission electron microscope operating at 30 kV using a STEM II detector in Field free mode.

To quantify the amount of cholesterol hybrid-g7-NPs-chol, NPs previously lyophilized to calculate the yield (~1 mg) were dissolved in chloroform (0.3 mL), followed by addition of isopropyl alcohol (0.6 mL) to precipitate the polymer. The mixture was vortexed (15 Hz for 1 min; ZX3, VelpScientifica, Usmate, Italy) and then filtered (polytetrafluoroethylene filter, porosity 0.20 μm, Sartorius). The amount of Chol in the sample was quantified by RP-HPLC using an HPLC apparatus comprised a Model PU980 pump provided with an injection valve with a 50 μL sample loop (Model 7725i Jasco) and an UV detector at 210 nm (UV975, Jasco). Chromatography separation was carried out on a Syncronics C18 (250×4.6 mm; porosity 5 μm; Thermo Fisher Scientific, Waltham, MA, USA) at r.t. and with a flow rate of 1.2 mL/min, operating in an isocratic mode using 50:50 v/v acetonitrile:ethanol as mobile phase. The solvents of the mobile phase were filtered through 0.45 μm hydrophilic polypropylene membrane filters (Sartorius) before their use. Chromatographic peak areas of the standard solutions were collected and used for the generation of calibration curves. Linearity was assumed in the range of 50-500 μg/mL (r2=0.99). All data are expressed as the mean of at least three determinations.

The chemico-physical properties, concentration and cholesterol amount in g7-NPs used in this work are described in table S1 along with morphological analysis with AFM and STEM (figure S1).

### 2.4 Colony management

All animal experiments were approved and carried out in accordance with Italian Governing Law (D.lgs 26/2014; Authorization n.581/2019-PR issued July 29, 2019 by Ministry of Health); the NIH Guide for the Care and Use of Laboratory Animals (2011 edition) and EU directives and guidelines (EEC Council Directive 2010/63/UE).

Our R6/2 colony lifespan was approximately of 12-13 weeks and only males were used to maintain it [22]. Transgenic 6-week-old R6/2 males were mated with wild type females (B6CBAF1/J, purchased from Charles River). CAG repeat length that could affect strain productivity, general behavior, litter size, pup survival, genotype frequency, phenotype was monitored every 6 months with a range between 150-180 CAGs.

Mice were weaned and then genotyped at 3 weeks of age (+/−3 days) and they were housed under standard conditions in enriched cage (22 ± 1°C, 60% relative humidity, 12 hours light/dark schedule, 3-4 mice/cage, with food and water ad libitum).

### 2.5 Mice treatments

For biodistribution studies, cholesterol release study and quantitative analysis, 7-week-old wt or R6/2 mice were treated with 1, 2 or 3 ip injections. For chronic experiments, R6/2 mice were treated with 2 ip injections at week, from 5 to 9 weeks of age. Wt and R6/2 littermates treated with saline solution was used as controls. In all the experiments, mice received 660 μg of cholesterol in each injection.

### 2.6 Immunohistochemistry and image acquisition

Animals were deeply anesthetized with Avertin 2.5% and transcardially perfused with PFA 4%. Brains, lungs and liver were collected in PFA 4% for 2h and then in 30% sucrose to prevent ice crystal damage during freezing in OCT. 15 μm-thick brain coronal sections or lung and liver sections were counterstained with the nuclear dye Hoechst 33258 (1:10.000, Invitrogen) and then mounted under cover slips using Vectashield (Vector Laboratories). To study the co-localization of g7-NPs-chol_2.0 with neuronal and glial markers, 15 μm-thick brain coronal sections were incubated with the following primary antibodies for 3h at RT: rabbit anti-DARPP32 (1:100, Cell Signalling, 2306); mouse anti-NeuN (1:100, Millipore, MAB377); rabbit anti-GFAP (1:250, Dako, Z0334); rabbit anti-IBA1 (1:100, Wako, 019-1971). Anti-rabbit Alexa Fluor 488-conjugated goat secondary antibodies (1:500; Invitrogen) or anti-mouse Alexa Fluor 488-conjugated goat secondary antibodies (1:500; Invitrogen) were used for detection (1h at RT). Sections were counterstained with the nuclear dye Hoechst 33258 (1:10.000, Invitrogen) and then mounted under cover slips using Vectashield (Vector Laboratories).

Images were acquired the following day with a confocal microscope (Leica SP5). Laser intensity and detector gain were maintained constant for all images and 3-z steps images were acquired at 40x. To quantify hybrid g7-NPs-chol in different tissues, ImageJ software was used to measure the fluorescence of Cy5 (n = 4 images/mouse/tissue).

To quantify the released bodipy cholesterol from g7-NPs-chol_2.0, Volocity software was used using the plug-in “find objects” and “calculate object correlation” (n = 6 images/mouse/tissue).

### 2.7 Liquid chromatography-mass spectrometry (LC-MS) analysis for d6-chol

A recently validated method was used [20]. Briefly, 50 μL of plasma was diluted with 200 μL of ethanol containing 200 ng of beta-sitosterol, used as internal standard. Samples were vortexed and centrifuged at 13 200 rpm for 15 min and aliquots of the supernatants were injected directly into the LC-MS system. Forty milligrams of each brain area and peripheral tissue were homogenized in 1 mL of ethanol/water 4:1 (v/v), containing 500 ng of internal standard. Homogenates were centrifuged for 15 min at 13200 rpm at 4 °C, and aliquots of the supernatants were injected into the LC-MS system. D6-chol levels were determined using a 1200 Series HPLC system (Agilent Technologies, Santa Clara, CA, U.S.A.) interfaced to an API 5500 triple quadrupole mass spectrometer (Sciex, Thornhill, Ontario, Canada). The mass spectrometer was equipped with an atmospheric pressure chemical ionization (APCI) source operating in positive ion and multiple reaction monitoring (MRM) mode to measure the product ions obtained in a collision cell from the protonated [M - 11 O|· ions of the analytes. The transitions identified during the optimization of the method were *m/z* 375.3-152.1 (quantification transition) and *m/z* 375.3-167.1 (qualification transition) for D6-chol; *m/z* 397.3–147.1 (quantification transition) and *m/z* 397.3-161.1 (qualification transition) for β-sitosterol (IS). D6-chol and beta-sitosterol were separated on a Gemini C18 column (50 × 2 mm; 5 μm particle size), using an isocratic gradient in 100 % methanol at 35 °C.

### 2.8 Gas chromatography-mass spectrometry (GC-MS) analysis for neutral sterols

To a screw-capped vial sealed with a Teflon-lined septum were added 50 μL of homogenates together with 500 ng of D4-lathosterol (CDN Isotopes, Canada), 500 ng of D6-desmosterol (Avantipolar Lipids, USA), and 100 ng of D6-lanosterol (Avantipolar Lipids, USA) as internal standards, 50 μL of butylated hydroxytoluene (BHT) (5 g/L) and 25 μL of EDTA (10 g/L). Argon was flushed through to remove air. Alkaline hydrolysis was allowed to proceed at room temperature (22°C) for 1h in the presence of 1 M ethanolic potassium hydroxide solution under magnetic stirring. After hydrolysis, the neutral sterols (lathosterol, desmosterol and lanosterol) were extracted twice with 5ml of hexane. The organic solvents were evaporated under a gentle stream of nitrogen and converted into trimethylsilyl ethers with BSTFA-1% TMCS (Cerilliant, USA) at 70 °C for 60 min. Analysis was performed by gas chromatography - mass spectrometry (GC-MS) on a Clarus 600 gas chromatograph (Perkin Elmer, USA) equipped with Elite-5MS capillary column (30 m, 0.32 mm, 0.25μm. Perkin Elmer, USA) connected to Clarus 600C mass spectrometer (Perkin Elmer, USA). The oven temperature program was as follows: initial temperature 180 °C was held for 1 min, followed by a linear ramp of 20 °C/min to 270 °C, and then a linear ramp of 5 °C/min to 290 °C, which was held for 10 min. Helium was used as carrier gas at a flow rate of 1 mL/min and 1 μL of sample was injected in splitless mode. Mass spectrometric data were acquired in selected ion monitoring mode. Peak integration was performed manually. Sterols were quantified against internal standards, using standard curves for the listed sterols.

### 2.9 Behavioral tests

Mice behavior was evaluated at 9 and 11 weeks of age.

Rotarod test: mice were tested over three consecutive days. Firstly, animals were trained on a rotating bar at 4 rpm for 5 minutes (apparatus model 47600, Ugo Basile). One hour later, mice were tested for three consecutive accelerating trials of 5 minutes with the rotarod speed linearly increasing from 4 to 40 rpm. The latency to fall from the rod was recorded for each trial and averaged.

Activity Cage test: animals were placed in an arena (25 cm × 25 cm) (2Biological Instrument) and allowed to freely move for an hour in presence of a low-intensity white light source. Movements were assessed by an automated tracking system (Actitrack software, 2Biological Instrument) connected to infrared sensors surrounding the arena. Total distance travelled, mean velocity speed, and numbers of rearings were analyzed. The % of time that mice explored the periphery or the center area of the was evaluated as a measure of anxiety-like behavior.

Novel Object Recognition (NOR) test: in the habituation stage, mice were placed into an empty non-reflective arena (44 × 44 × 44 cm) for 10 minutes. In the familiarization stage, two identical objects (A’ and A”) were presented to each animal for 10 minutes. The day after, during the test stage, animals were exposed to one familiar object (A’) and a new object (B) for 10 minutes. All phases of the test were conducted with a low-intensity white light source. The index of discrimination was calculated as (time exploring the novel object-time exploring the familiar object)/(time exploring both objects) × 100. Mice exploring less than 7 sec. were excluded from the analysis due to their inability to perform the task.

Paw clasping test: animals were suspended by the tail for 30 seconds and the clasping phenotype was graded according to the following scale: level 0, no clasping; level 1, clasping of the forelimbs only or both fore- and hindlimbs once or twice; and level 2, clasping of both fore- and hindlimbs more than three times or more than 5 s.

Grip strength test: animals were lifted by the tail, lowered towards the grip (Ugo Basile) and gently pulled straight back with consistent force until they released its grip. The forelimb grip force, measured in grams, was recorded. The test was repeated for 5 times, and measures were averaged.

### 2.10 Bio-Plex

Animals were deeply anesthetized with Avertin 2.5% to collect blood which was centrifuged at 13.000 rpm at 4°C for 15 minutes to obtain the plasma. Striatum, cortex, and liver were isolated and frozen. 10 mg of striatum, cortex and liver were homogenize using a tissue grinder in 1 mL of lysing solution according to manufacturer instructions (Bio-Plex^®^ Cell Lysis Kit, Biorad, #171304011). The lysate was frozen at −80°C, sonicated at 40% for 20 seconds and centrifuged at 4.500 *rcf* at 4°C for 4 minutes to collect the supernatant. The supernatant was quantified using DC™ Protein Assay Kit I (Biorad, #5000111) and samples were diluted to a final concentration of 500 μg/mL. To perform the Bio-Plex assay, 150 μL of assay buffer were added to 150 μL of samples.

Concerning the plasma, samples were centrifuged at 1.500 rcf at 4°C for 5 minutes. 60 μL of assay buffer and 120 μL of sample diluent were added to 60 μL of plasma.

All samples were tested for the following cytokines using the Bio-Plex Pro Mouse Cytokine 23-plex Assay: IL-1α, IL-1β, IL-2, IL-3, IL-4, IL-5, IL-6, IL-9, IL-10, IL-12 (p40), IL-12 (p70), IL-13, IL-17A, Eotaxin, G-CSF, GM-CSF, IFN-γ, KC, MCP-1 (MCAF), MIP-1α, MIP-1β, RANTES, TNF-α, (Biorad, #M60009RDPD) according to manufacturer instructions and detected using Bioplex™ 200 System (Bio-Rad). The concentration of each cytokine was calculated through calibration curve (individual for each cytokine), determined independently for each experiment, by Bioplex Manager^TM^ software 4.1.

### 2.11 Statistics

Prism 8 (GraphPad software) was used to perform statistical analyses. G-power software was used to pre-determine group allocation, data collection and all related analyses. For animal studies, mice were assigned randomly, and sex was balanced in the various experimental groups; animals from the same litter were divided in different experimental groups; blinding of the investigator was applied to *in vivo* procedures and all data collection. Grubbs’ test was applied to identify outliers. For each set of data to be compared, we determined whether data were normally distributed or not to select parametric or not parametric statistical tests. The specific statistical test used is indicated in the legend of all results figures. **Table S3** summarizes all the trials and read-outs performed.

## 3. Results and discussion

### 3.1 Localization of hybrid-g7-NPs-chol MIX-N and MIX-SE *in vivo*

The characteristics of both hybrid-g7-NPs-chol MIX-N and MIX-SE were in line with those described in previous studies [13, 21] in terms of size, homogeneity, surface charge, cholesterol content, and morphology (Table S1 and Fig. S1).

The hybrid-g7-NPs-chol MIX-N and MIX-SE were first tested in 7-week-old wild-type (wt) mice to verify their uptake and distribution *in vivo.* To this aim, animals were treated with a single or multiple (3) intraperitoneal (ip) injections of hybrid-g7-NPs-chol MIX-N or MIX-SE labeled with cyanine 5 (Cy5) and sacrificed at different time points (4 h, 24 h, 1 week, 2 weeks) following the last ip injection (Fig. 1(a)). Fluorescence analysis was then performed on brain and peripheral slices to analyze the localization of the Cy5-labeled hybrid-NPs-Chol signals of both MIXs (Fig. 1(b) and 1(c); Fig. S2). Four hours after a single ip injection, Cy5 signal was detected in the striatum, cortex (Fig. 1(b) and 1(c)), and hippocampus, as well as in lung and liver (Fig. 1(b) and 1(c) and Fig. S2(a) and S2(b)), indicating that BBB transit was rapid. Quantification of Cy5 signals revealed that the kinetics of both MIXs in peripheral tissues were quite rapid, at least in lung and somewhat less so in liver, since the Cy5 signals decreased markedly 24 h after the ip injection (Fig. 1(b) and 1(c); Fig. S2). Importantly, NPs were accumulating in the brain over time, as evidenced by the Cy5-NPs signal being present at 24 h, 1 week, and 2 weeks after ip injection (Fig. 1(b) and 1(c); Fig. S2), and strengthening following multiple injections (Fig. 1(b) and 1(c); Fig. S2). High-magnification confocal images indicated the presence of hybrid-g7-NPs-chol in different neuronal and glial cell types as demonstrated by the colocalization of the signals from Cy5 (NPs) and DARPP32 (marker of striatal medium spiny neurons), GFAP (marker of astrocytes), and IBA1 (marker of microglia) (Fig. 1(d)). Although the biodistributions of MIX-N and MIX-SE were similar, the latter showed higher aggregation in liver, a finding that was more evident after multiple ip injections but which disappeared after 1 week (Fig. S2(b)).

**Figure 1.**
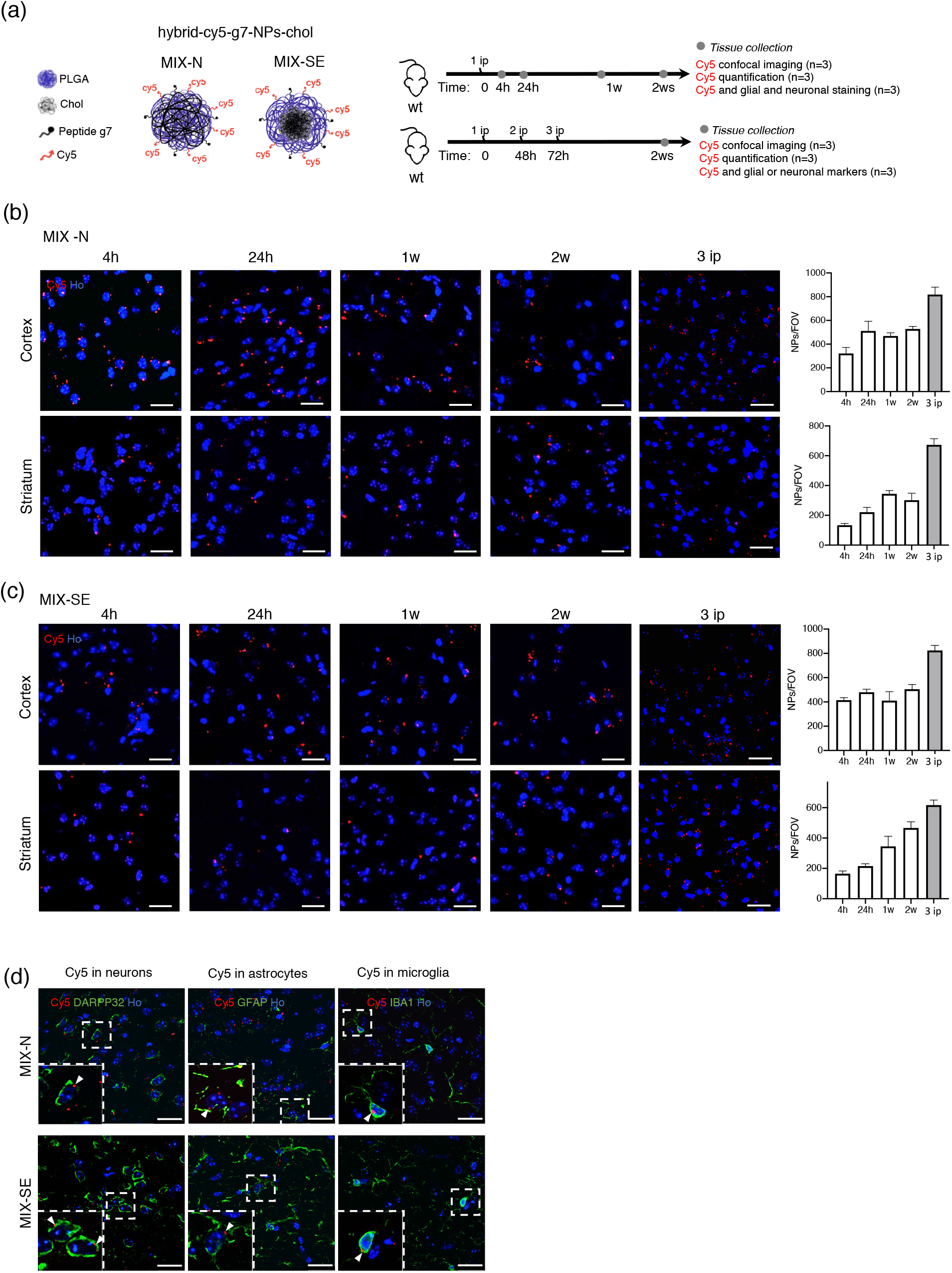
*In vivo* distribution of hybrid-cy5-g7-NPs-chol. (a) Experimental paradigm used in the study. Wild-type (wt) mice (n = 2 mice/MIX/time point) were treated with a single or multiple intraperitoneal (ip) injection of hybrid-Cy5-g7-NPs-chol (MIX-N or MIX-SE) and sacrificed at different time points. Brain, liver, and lung were collected for distribution analysis. (b) and (c) Representative confocal images of coronal slices containing cortex and striatum from wt mice that received 1 or 3 ip injections of hybrid-Cy5-g7-NPs-chol (MIX-N in (b) or MIX-SE in (c)) and sacrificed after 4 h, 24 h, 1 week, and 2 weeks with relative quantification. (d) Representative confocal images of immunostaining for DARPP32, GFAP, and IBA1 (green) on coronal sections of brains isolated from wt mice after receiving ip-injected hybrid-Cy5-g7-NPs-chol labeled with Cy5 (red) and sacrificed 2 weeks after the injection. White arrowheads indicate intracellular g7-NPs. Hoechst was used to counterstain nuclei (Ho, blue). Scale bar is 30 μm. Data are expressed as the number of g7-NPs per field of view ± standard error of the mean.

Overall, these results demonstrate that hybrid-g7-NPs-chol rapidly reach the brain and accumulate over time within target cells.

### 3.2 Cholesterol delivery and intracellular release

To track cholesterol delivery and intracellular release from hybrid-g7-NPs-chol, we performed specific experiments using g7-NPs-chol covalently labeled with Cy5 and loaded with the fluorescent analogue bodipy cholesterol (hybrid-Cy5-g7-NPs-bodipy-chol). Seven-week-old wt and HD mice (R6/2 model) [22] were treated with a single ip injection of hybrid-Cy5-g7-NPs-bodipy-chol MIX-N or MIX-SE and sacrificed at 24 h and 2 weeks after the ip injection (Fig. 2(a)). Following confocal analysis in brain slices, we analyzed the colocalization of red spots (Cy5, NPs) and the green signal (bodipy cholesterol) to evaluate bodipy cholesterol release from the new formulations. We show that 24 h after a single ip injection of hybrid-Cy5-g7-NPs-bodipy-chol, Cy5 and bodipy-chol signals nicely colocalized in striatum and cortex, as indicated by the scatterplot of red and green pixel intensities (Fig. 2(b) and 2(c)). Importantly, analysis performed 2 weeks after ip injection revealed a partial separation between Cy5 and bodipy-chol signals (Fig. 2(b) and 2(c)), indicating a slow and progressive release of cholesterol over time. In contrast, cholesterol release in the liver of mice treated with MIX-N was faster than in the brain. In fact, a complete overlap of red and green signal was found 24 h after the ip injection, while 2 weeks after the ip injection all the exogenous cholesterol was released from NPs (Fig. S3). By comparing the overlap coefficient between Cy5 and bodipy signals (Table S2), we found that approximately 30% of bodipy-chol no longer colocalized with Cy5-NPs in cortex and striatum starting from 2 weeks after ip injection, suggesting a progressive release from hybrid-g7-NPs-chol in the brain, in parallel with a reduction in Cy5 signal, probably due to polymer degradation. In contrast, about 90% of bodipy-cholesterol no longer colocalized with Cy5-NPs in the liver (Table S2). No differences in cholesterol release kinetics were found between wt and HD mice, indicating that cholesterol release over-time did not depend on mouse genotype (Table S2). We also conclude that MIX-N and MIX-SE NPs had similar biodistribution profiles and kinetics of cholesterol release. Notably, as MIX-N showed less aggregation in liver and, more importantly, as the surfactant present in the formulation (Pluronic F68) is approved by the FDA [23], we decided to proceed by testing only this kind of NPs in subsequent studies.

**Figure 2.**
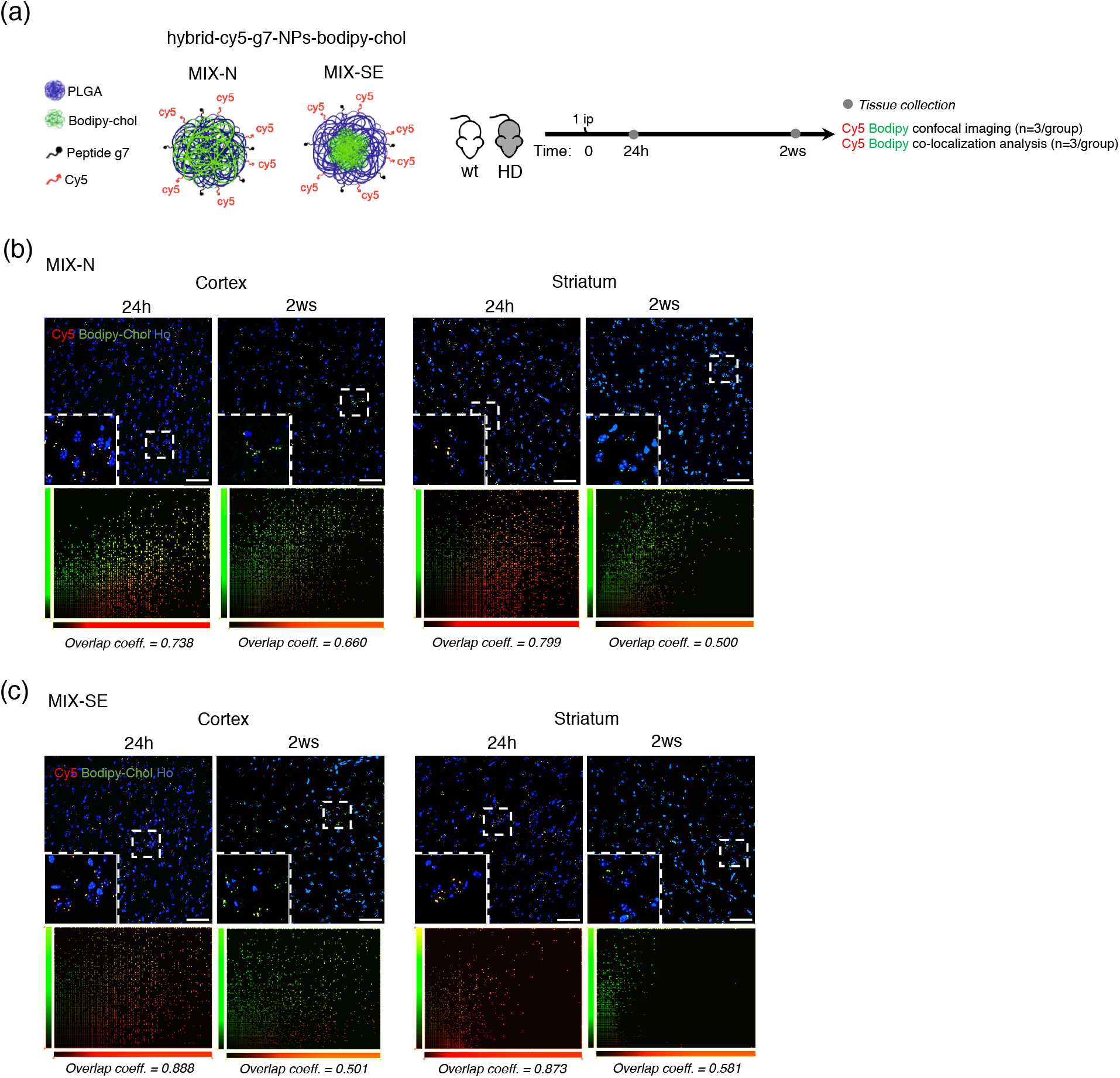
Cholesterol release from hybrid-cy5-g7-NPs-bodipy-chol: qualitative analysis. (a) Experimental paradigm used in the study. Wild-type and R6/2 mice (n = 3 mice/genotype/MIX/time point) were treated with a single ip injection of hybrid-Cy5-g7-NPs-bodipy-chol (MIX-N or MIX-SE) and sacrificed at different time points. Brain and liver were collected for the analysis. (b) and (c) Representative confocal images of brain slices from R6/2 mice after ip injection of hybrid-Cy5-g7-NPs-bodipy-chol (MIX-N in (b) or MIX-SE in (c)) and sacrificed after 24 h or 2 weeks and relative co-localization of bodipy-chol and g7-NPs. Hoechst was used to counterstain nuclei (Ho, blue). Scale bar is 50 μm.

### 3.3 Kinetics of cholesterol delivery to different organs

Next, to study the kinetics of exogenous cholesterol delivery and to quantify the total amount of cholesterol delivered *in vivo*, we performed two experiments using only MIX-N hybrid-g7-NPs-chol loaded with cholesterol labeled with six deuterium atoms (d6-chol) (hybrid-g7-NPs-d6-chol).

In the first experiment, 7-week-old HD mice were treated with a single ip injection of hybrid-g7-NPs-d6-chol and sacrificed at 30 min, 6 h, 24 h, 1 week, and 2 weeks after the ip injection (Fig. 3(a)). Blood, kidney, lung, liver, cortex, striatum, and cerebellum were collected to measure the amount of d6-chol in the different tissues using liquid chromatography mass spectrometry (LC-MS), starting from 6 h after the ip injection (Fig. 3(b)). We show that starting from 24 h after the treatment, the content of d6-chol increased in the tissues over a 2-week period, a finding which is indicative of its slow and progressive release from the NPs (Fig. 3(b)). In contrast, in the liver, d6-chol was rapidly released 30 min after ip injection and rapidly degraded over time. In lung and kidney, NPs were detected around 6 h after the ip injection, the peak of cholesterol release occurred 1 week after the ip injection, after which it was rapidly eliminated (Fig. 3(c)). In plasma, the maximum amount of cholesterol was detected 24 h after the ip injection (Fig. 3(c)). Importantly, this concentration is not significant compared with the amount of cholesterol that is present in mouse blood (128 mg/100 mL).

**Figure 3.**
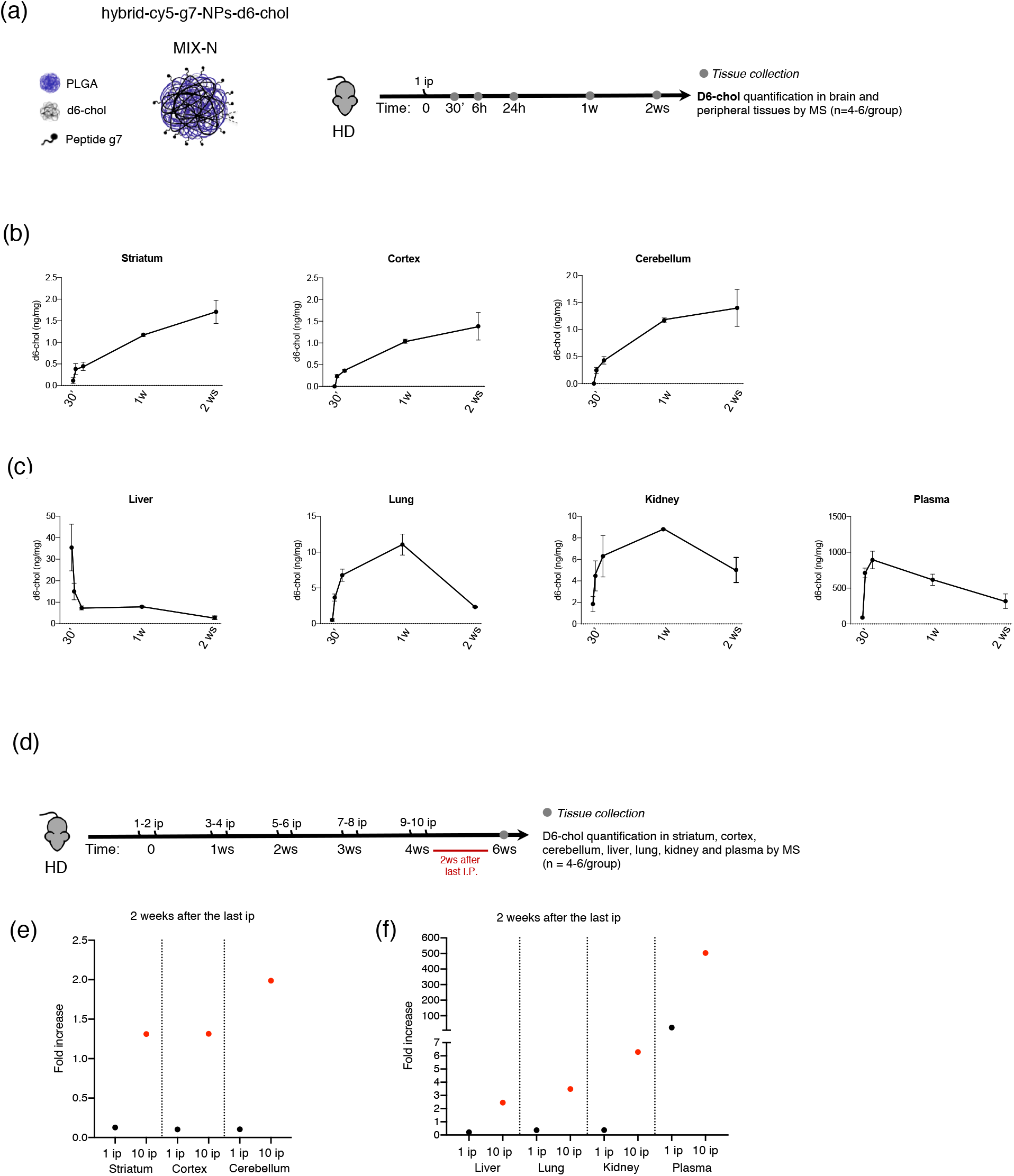
Cholesterol release from hybrid-cy5-g7-NPs-d6-chol: quantitative analysis. (a) Experimental paradigm used in the study. R6/2 mice were treated with a single ip injection of hybrid-g7-NPs-d6-chol (MIX-N) and sacrificed at different time points. Striatum, cortex, cerebellum, liver, lung, kidney, plasma, and liver were collected for mass spectrometry analysis (n = 3 mice/time point). (b) and (c) Levels of d6-chol in striatum, cortex, cerebellum (b), liver, lung, kidney, and plasma (c) measured by LC-MS. (d) Experimental paradigm used in the study. R6/2 mice were treated with hybrid-g7-NPs-d6-chol (MIX-N) from 5 weeks of age to 9 weeks of age with 2 ip injections/week and sacrificed 2 weeks after the last ip injection. Striatum, cortex, cerebellum, liver, lung, kidney, and plasma and liver were collected for mass spectrometry analysis (n = 3 mice). (e) and (f) Levels of d6-chol in striatum, cortex, cerebellum (e), liver, lung, kidney, and plasma (f) measured by LC-MS (red dots). Black dots refer to the measurement represented in figure 3B-C. Data are expressed as means ± standard error of the mean.

In a second experiment, to quantify cholesterol delivered with the therapeutic regimen of interest, HD mice were treated from 5 to 9 weeks of age with 2 ip injections/week and sacrificed 2 weeks after the last ip injection (Fig. 3(d)). MS analysis revealed that the concentration of d6-chol measured in each tissue following 10 ip injections was around 10 times the concentration of d6-chol measured 2 weeks after a single ip injection (Fig. 3(d) and (e)), indicating that the exogenous cholesterol accumulated in all tissues, even if with different kinetics.

These results demonstrated that the kinetics of cholesterol release differed between brain, plasma, and peripheral tissues and that this delivery system allows a slow release and accumulation of cholesterol in different brain regions where it becomes available to cells over time. Moreover, the fast elimination of cholesterol from blood and peripheral tissues potentially avoids systemic side effects after chronic treatment. Finally, these data combined with the data obtained with bodipy-chol (Fig. 2(b) and 2(c)), support furtherly the hypothesis that the release of cholesterol from NPs is progressive and slow.

### 3.4 Enhancement of cholesterol synthesis in the HD mouse brain

We have recently demonstrated that a therapeutically relevant dose of cholesterol delivered to the striatum through osmotic mini-pumps indirectly stimulated endogenous cholesterol synthesis, leading to the reversal of both motor and cognitive abnormalities in HD mice [19]. In contrast, lower doses delivered to the striatum through osmotic mini-pumps [19] or to the brain via PLGA-g7-NPs-chol [13] led to a complete rescue of cognitive decline without significant change in endogenous brain cholesterol biosynthesis or in motor performance.

Since hybrid-g7-NPs-chol [21], with their hybrid structure, are able to carry a larger amount of cholesterol than the previously used PLGA-g7-NPs-chol [13], we sought to test whether the increased amount of cholesterol in hybrid-g7-NPs-chol was sufficient to stimulate endogenous cholesterol synthesis in the diseased brain. As surrogate markers of cholesterol biosynthesis, we quantified cholesterol precursors (lanosterol, lathosterol, desmosterol) by isotopic dilution gas-chromatography mass spectrometry (ID-MS) in the striata of HD mice after a chronic treatment. Accordingly, HD mice from 5 to 9 weeks of age were treated with PLGA-g7-NPs-chol or with hybrid-g7-NPs-chol with 2 ip injections/week and sacrificed 2 weeks after the last ip injection (Fig. 4(a)). As expected, robust deficits of lanosterol, lathosterol, and desmosterol were evident in striatum from HD mice treated with saline compared with wt littermates (Fig. 4(b)-(d)), confirming previous results [15–18]. Of note, significant increases in lanosterol and desmosterol levels were found in striatal tissues of HD mice treated with hybrid-g7-NPs-chol compared with those treated with PLGA-g7-NPs-chol (Fig. 4(b)-(d)).

**Figure 4.**
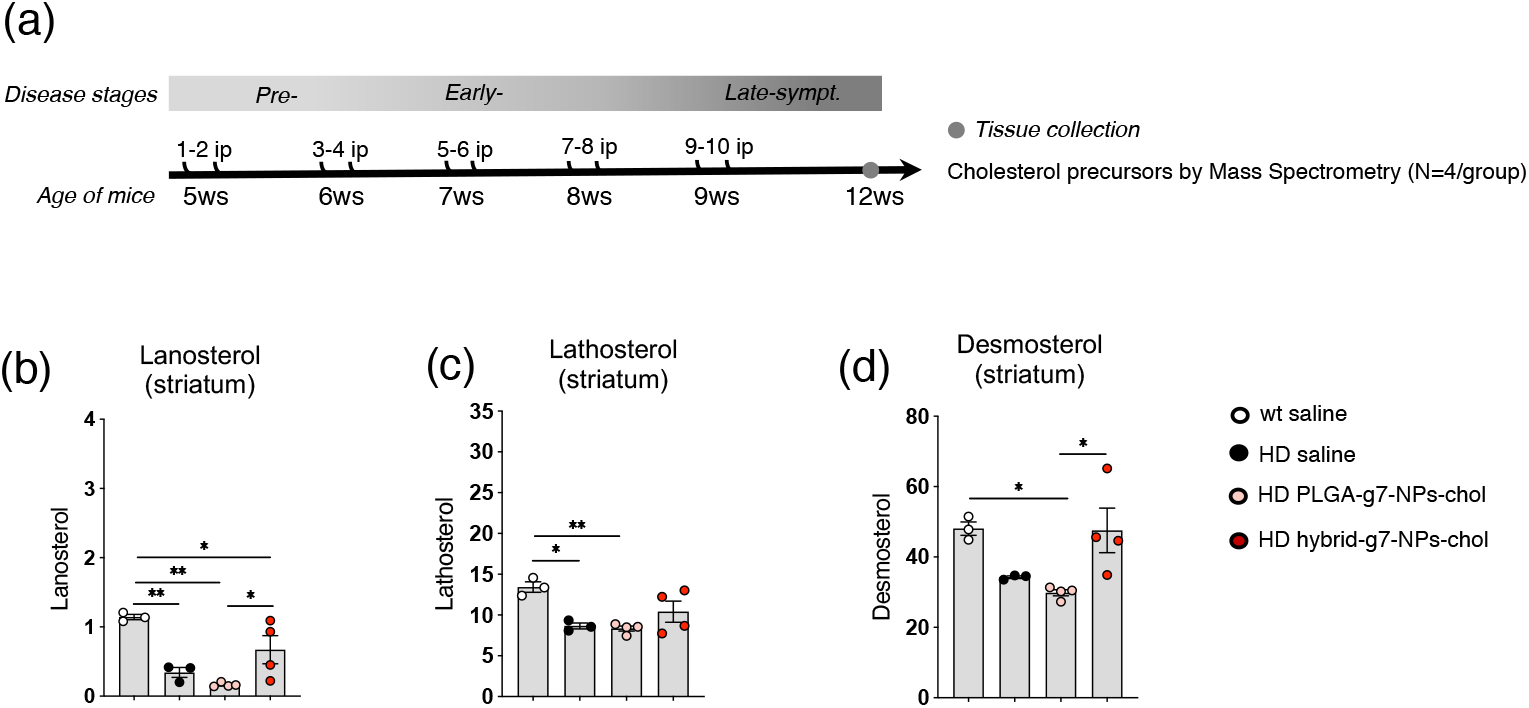
Endogenous cholesterol synthesis following systemic and chronic injection of hybrid g7-NPs-chol. (a) Experimental paradigm used in the study. R6/2 mice were treated with PLGA-g7-NPs-chol and hybrid-g7-NPs-chol from 5 weeks of age to 9 weeks of age with 2 ip injections/week. Wt and R6/2 littermates were treated with saline solution as controls. Striatum and cortex were collected at 11 weeks of age for mass spectrometry analysis (n = 3-4 mice/group). (b), (c), and (d): Lanosterol (b), lathosterol (c), and desmosterol (d) levels measured by GC-MS in the striatum of wt saline, R6/2 saline, R6/2 + PLGA-g7-NPs-chol, and R6/2 + hybrid-g7-NPs-chol mice at 11 weeks of age (n = 3–4 mice/group). Data are expressed as means ± standard error of the mean. Each dot corresponds to the value obtained from each animal. Statistics: one-way ANOVA with Newman-Keuls post-hoc test (*p<0.05; **p<0.01; ***p<0.001).

Taken together, these results suggest that the hybrid-g7-NPs-chol transport and release in the brain more cholesterol compared with PLGA-g7-NPs-chol and that the dose is able to enhance endogenous cholesterol biosynthesis in HD mice.

### 3.5 Effects on cognition, locomotion, and strength

To assess the power of hybrid-g7-NPs-chol to counteract motor and cognitive defects in HD mice, hybrid-g7-NPs-chol were ip injected into R6/2 mice with the same experimental paradigm described in Fig. 4(a) and their motor and cognitive performance were compared with those of R6/2 and wt mice treated with saline solution.

First, we analyzed motor coordination by evaluating the latency of mice to fall when tested on a rotating bar with accelerating speed in the rotarod test. Starting from 8 weeks of age, HD mice exhibited a progressive deterioration in motor coordination, as shown by the shorter latency to fall compared with wt controls. Systemic and chronic administration of hybrid-g7-NPs-chol did not rescue this defect in HD mice (Fig. 5(a)).

**Figure 5.**
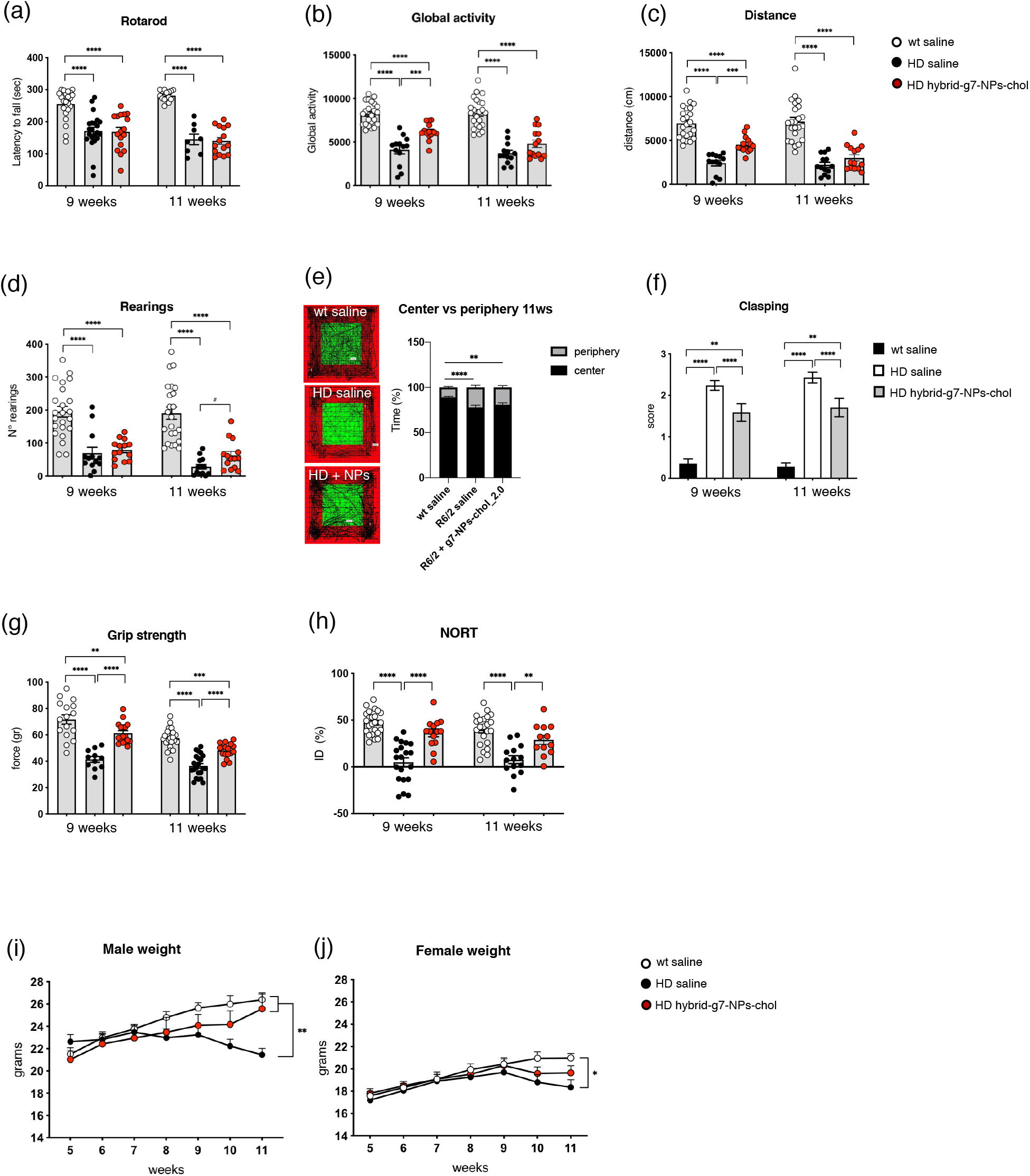
Cognitive and motor abilities of HD mice following systemic and chronic injection of hybrid g7-NPs-chol. (a) Latency to falling (seconds) from an accelerating rotarod at 8 and 10 weeks of age in wt saline (n = 24-25), R6/2 saline (n = 15-17), and R6/2 + hybrid-g7-NPs-chol (n = 17) treated mice. (b), (c), (d), (e) and (f): Global motor activity (b), total distance traveled (c), and number of rearings (d) in an open-field test at 9 and 11 weeks of age in wt saline (n = 24-25), R6/2 saline (n = 15-17), and R6/2 + hybrid-g7-NPs-chol (n = 17) mice. (e) Representative track plots generated from the open-field test from wt saline (n = 24-25), R6/2 saline (n = 15-17), and R6/2 + hybrid-g7-NPs-chol (n = 17) mice and relative quantification of the times spent in the center and in the periphery (%) of the arena at 11 weeks of age. (f) Grip strength (grams) at 9 and 11 weeks of age in wt saline (n = 16-24), R6/2 saline (n = 11-22), and R6/2 + hybrid-g7-NPs-chol (n = 17) mice. (g) Paw clasping at 9 and 11 weeks of age in wt saline (n = 25), R6/2 saline (n = 22), and R6/2 + hybrid-g7-NPs-chol (n = 17) mice. (h) Discrimination index (DI %) in the novel object recognition test of wt saline (n = 22-23), R6/2 saline (n = 13-19), and R6/2 + hybrid-g7-NPs-chol (n = 10-13) mice at 9 and 11 weeks of age. DI above zero indicates a preference for the novel object; DI below zero indicates a preference for the familiar object. (i) and (j) Body weight (grams) of male (i) and female (j) mice at different time points. Data are from three independent trials and shown as scatterplot graphs with means ± standard error. Each dot corresponds to the value obtained from each animal. Statistics: one-way ANOVA with Newman-Keuls post-hoc test (*p<0.05; **p<0.01; ****p<0.0001; ***p<0.001) or unpaired Student t-test (#p<0.05; ##p<0.01).

To further test the animals’ motor abilities, we analyzed spontaneous locomotory activity in the activity cage test. During disease progression, HD mice showed a severe hypokinetic phenotype as demonstrated by reduced global activity, total distance travelled, and number of rearings compared with wt mice, at both 9 and 11 weeks of age. At 9 weeks, HD mice treated with hybrid-g7-NPs-chol had greater global activity and total distance travelled compared with HD mice treated with saline, even if they did not reach the performance observed in wt mice. Moreover, these differences were lost at 11 weeks of age, suggesting that the amount of cholesterol delivered in the brain was not sufficient to counteract motor deficits at a late symptomatic time point, when the HD phenotype worsens (Fig. 5(b) and (c)). When we looked at the number of rearings, no rescue was measured in HD mice treated with hybrid-g7-NPs-chol (Fig. 5(d)). As a measure of anxiety-like behavior, the time that mice spent exploring the periphery or center area of the arena during the activity cage test was also evaluated (Fig. 5(e)). HD animals spent more time in the periphery compared with wt mice, indicating anxiety-related behavior. Cholesterol delivery did not rescue this phenotype (Fig. 5(e)).

As a marker of disease progression, we measured hind-limb clasping with the paw clasping test, a test widely used to measure neurological features in several mouse models of neurodegeneration. In HD mice treated with hybrid-g7-NPs-chol, this phenotype was ameliorated (Fig. 5(f)).

To study neuromuscular functions and strength, we determined the force developed by the mice using the grip-strength test. Muscular strength was reduced in HD mice from 9 weeks of age and it was completely rescued by hybrid-g7-NPs-chol at both 9 and 11 weeks of age (Fig. 5(g)). Furthermore, we found that 44% of the analyzed R6/2 animals treated with saline suffered from epileptic seizures, while only 18% of R6/2 mice injected with hybrid-g7-NPs-chol were affected. Finally, to evaluate cognitive function we performed the novel object recognition (NOR) test. As expected, long-term memory declined during disease progression in HD mice, with a marked impairment in the ability to discriminate novel and familiar objects at 11 weeks of age. HD mice treated with hybrid-g7-NPs-chol performed similarly to wt mice, indicating that this treatment completely prevented cognitive decline in these animals (Fig. 5(h)). Weight loss was observed in R6/2 mice starting from a late time point (10 weeks of age). Remarkably, this parameter was rescued in male R6/2 mice treated with hybrid-g7-NPs-chol (Fig. 5(i) and 5(j)).

Collectively, these results indicate that the dose of cholesterol delivered and released in the brain with chronic treatment was sufficient to prevent cognitive decline over time and ameliorate some motor defects at 9 weeks of age. However, the fast and aggressive phenotype of this HD mouse model did not allow us to evaluate the long-term effect of this treatment when all cholesterol is released from the NPs, which may require several weeks.

### 3.6 Assessment of markers of inflammation, a possible side effect

To explore any eventual side effects of chronic administration of hybrid-g7-NPs-chol, we next sought to analyze the inflammation status of treated mice. Cytokines, chemokines, and growth factors are cell-signaling proteins that mediate a wide range of physiological responses including immunity and inflammation, and are also associated with a spectrum of neurodegenerative diseases [24–25]. Through the simultaneous detection of 23 analytes in a single well of a 96-well microplate, we analyzed the inflammation status of striatum, cortex, liver, and plasma from HD mice treated with saline or with hybrid-g7-NPs-chol. In general, we did not observe gross changes in the levels of the analytes analyzed, except for an increase in IL-2 in the striatum and a decrease in eotaxin in cortex and in IL-1a and IL2 in plasma of R6/2 mice treated with hybrid-g7-NPs-chol compared with R6/2 mice treated with saline (Table 1). These findings suggest that chronic treatment with hybrid-g7-NPs-chol is safe in R6/2 mice. Moreover, observation of the mice during chronic administration regimens did not reveal any cases of mortality in the treated and control groups and no signs of abnormal behavioral reactions and general clinical symptoms were detected.

**Table 1.** Inflammatory response of HD mice following systemic and chronic injection of hybrid g7-NPs-cho. Analysis of cytokines and chemokines by Bio-Plex array on the striatum, cortex, liver, and plasma of HD mice treated with saline solution vs. HD mice treated with hybrid-g7-NPs-chol (n = 4–5 mice/group). Data represent the fold-change and are expressed as means ± standard error of the mean. Statistics: unpaired Student t-test (*p<0.05).

| Analyte | Striatum |  | Cortex |  | Liver |  | Plasma |  |
| --- | --- | --- | --- | --- | --- | --- | --- | --- |
|  | fold-change |  | fold-change |  | fold-change |  | fold-change |  |
| | Mean ± SEM | p-value | Mean ± SEM | p-value | Mean ± SEM | p-value | Mean ± SEM | p-value |
| IL-1a | 1,010 ± 0,025 | 0,9348 | 0,927 ± 0,033 | 0,3884 | 0,976 ± 0,111 | 0,8622 | **0,212 ± 0,032** | **0,0078 \*\*** |
| IL-1b | 1,055 ± 0,200 | 0,8435 | 0,747 ± 0,134 | 0,3254 | 1,088 ± 0,184 | 0,7466 | 0,920 ± 0,381 | 0,8608 |
| IL-2 | **1,415 ± 0,076** | **0,0081 \*\*** | 0,708 ± 0,013 | 0,1058 | 1,159 ± 0,179 | 0,4831 | **0,398 ± 0,081** | **0,0467 \*** |
| IL-3 | 1,012 ± 0,298 | 0,9739 | 0,771 ± 0,097 | 0,2840 | 1,031 ± 0,038 | 0,7151 | N/D | |
| IL-4 | 0,632 ± 0,110 | 0,5871 | 1,472 ± 0,499 | 0,4103 | 0,744 ± 0,182 | 0,4333 | 0,429 ± 0,184 | 0,3575 |
| IL-5 | 1,264 ± 0,165 | 0,5834 | 1,276 ± 0,561 | 0,6831 | 0,924 ± 0,158 | 0,7211 | 0,330 ± 0,085 | 0,1610 |
| IL-6 | 1,061 ± 0,132 | 0,7723 | 1,019 ± 0,102 | 0,9132 | 0,889 ± 0,133 | 0,5446 | N/D | |
| IL-9 | 0,789 ± 0,143 | 0,4046 | 0,836 ± 0,145 | 0,5136 | 0,900 ± 0,112 | 0,6095 | N/D | |
| IL-10 | 1,020 ± 0,291 | 0,9593 | 0,677 ± 0,123 | 0,1301 | 1,109 ± 0,354 | 0,8100 | 1,175 ± 0,002 | 0,1610 |
| IL-12 (p40) | 1,666 ± 0,435 | 0,2450 | 0,667 ± 0,176 | 0,3861 | 1,843 ± 0,113 | 0,1025 | 0,921 ± 0,236 | 0,8124 |
| IL-12 (p70) | 0,944 ± 0,242 | 0,8601 | 0,798 ± 0,138 | 0,3859 | 0,803 ± 0,116 | 0,4620 | N/D | |
| IL-13 | 0,950 ± 0,072 | 0,5667 | 0,835 ± 0,069 | 0,2902 | 0,977 ± 0,097 | 0,8460 | 1,099 ± 0,152 | 0,5886 |
| IL-17 | 1,345 ± 0,414 | 0,5198 | 0,892 ± 0,125 | 0,2902 | 1,194 ± 0,104 | 0,3108 | 1,345 ± 0,414 | 0,5198 |
| Eotaxin | 1,695 ± 0,359 | 0,1447 | **0,447 ± 0,078** | **0,0228 \*** | 0,958 ± 0,215 | 0,8927 | 0,809 ± 0,195 | 0,5637 |
| G-CSF | 0,943 ± 0,203 | 0,8490 | 0,914 ± 0,153 | 0,6571 | 1,001 ± 0,120 | 0,9970 | 0,987 ± 0,148 | 0,9529 |
| GM-CSF | 1,025 ± 0,222 | 0,9314 | 1,060 ± 0,220 | 0,8302 | 1,090 ± 0,080 | 0,4218 | 0,724 ± 0,188 | 0,2956 |
| ING-gamma | 1,413 ± 0,272 | 0,2607 | 0,899 ± 0,102 | 0,5404 | 1,155 ± 0,184 | 0,4968 | 0,749 ± 0,322 | 0,7225 |
| KC | 1,337 ± 0,302 | 0,3446 | 1,340 ± 0,169 | 0,1438 | 0,988 ± 0,313 | 0,9742 | 0,489 ± 0,086 | 0,1494 |
| MCP-1 | 0,947 ± 0,375 | 0,9216 | 0,907 ± 0,196 | 0,8058 | 1,657 ± 0,414 | 0,2082 | 1,365 ± 0,689 | 0,7636 |
| MIP-1a | 1,133 ± 0,326 | 0,7593 | 0,816 ± 0,138 | 0,4708 | 1,065 ± 0,126 | 0,6874 | 0,525 ± 0,222 | 0,5987 |
| MIP-1b | 1,511 ± 0,186 | 0,1488 | 0,731 ± 0,137 | 0,3533 | 0,920 ± 0,192 | 0,7806 | 0,614 ± 0,452 | 0,4762 |
| Ranteg | 1,010 ± 0,075 | 0,9520 | 1,020 ± 0,134 | 0,9271 | 0,882 ± 0,087 | 0,5915 | 0,998 ± 0,231 | 0,9954 |
| TNF-a | 1,173 ± 0,276 | 0,6227 | 0,854 ± 0,249 | 0,6447 | 1,100 ± 0,352 | 0,8135 | 0,851 ± 0,371 | 0,8106 |

data expressed as fold-change (HD mice treated with hybrid-g7-NPs-chol vs HD mice treated with saline solution)

Overall, these results suggest that chronic administration of hybrid-g7-NPs-chol does not lead to side effects in HD mice.

## 4. Conclusion

Previous studies pointed out the benefits of strategies aimed at delivering cholesterol to the HD brain [13, 19], but defining the dose of cholesterol that reaches the brain is critical to a complete understanding of the power and limits of the approach. With the aim of developing a new and non-invasive strategy closer to clinical application, hybrid-g7-NPs-chol were produced with improved chemical and physical properties and increased cholesterol content [21]. Here, we characterized hybrid-g7-NPs-chol *in vivo* in a transgenic mouse model of HD. We demonstrated that hybrid-g7-NPs-chol are taken up and reach different cell types in the brain, and that they accumulate over time and are able to release cholesterol, which becomes available for neuronal functions. Importantly, NPs are rapidly degraded in the plasma and in peripheral tissues without a detectable inflammatory response. These systems can be optimized further in order to transport not only cholesterol but other molecules that can be useful in treating HD and other brain pathologies, or even used with different routes of administration [20]. Finally, we highlighted the utility of cholesterol as a model drug with which to define delivery systems based on NPs.

## Acknowledgments

The authors acknowledge the technical assistance of Dr. Chiara Cordiglieri, responsible of the INGM Imaging Facility (Istituto Nazionale Genetica Molecolare - INGM, Milan, Italy) and Centro Interdipartimentale Grandi Strumenti UNIMORE for AFM and SEM-FEG images. This research was supported by Telethon Foundation (GGP17102) to E.C., by Italian Ministry of Health (RF-2016-02361928) to M.S. and E.C., by MAECI Progetti di Grande Rilevanza Scientifica Italy-USA (MAE00691612020-06-26), PORFESR Emilia Romagna “Mat2Rep”, IMI2 grant “IM2PACT” to G.T., and Fondazione Umberto Veronesi Fellowship Grant to J.T.D.

## Author contributions

M.V., G.T., B.R. and E.C. conceived the study; G.B. performed immunostaining experiments and provided confocal images and quantification; G.B. and M.V. performed *in vivo* experiments; G.T., B.R., F.F. and M.A.V. conceptualized and developed NPs; J.T.D. and I.O. produced and characterized chemico-physical features of NPs. C.C., V.L. and F.T. performed GC-MS analysis and analyzed the data; A.P., M.F., L.C., R.B. and M.S. performed LC-MS analysis for d6-chol and analyzed the data; G.B., M.V. and M.B. performed bioplex analysis; M.V. and G.B. collected study data and performed statistical analyses; M.V. and E.C. oversaw and coordinated responsibility for all research activities and their performance and provided experimental advice throughout the work; E.C. secured the funding, the collaborations and the execution of the entire project. M.V., G.B., and E.C. wrote the paper that has been edited and reviewed by all authors.

---

**Figure S1.**
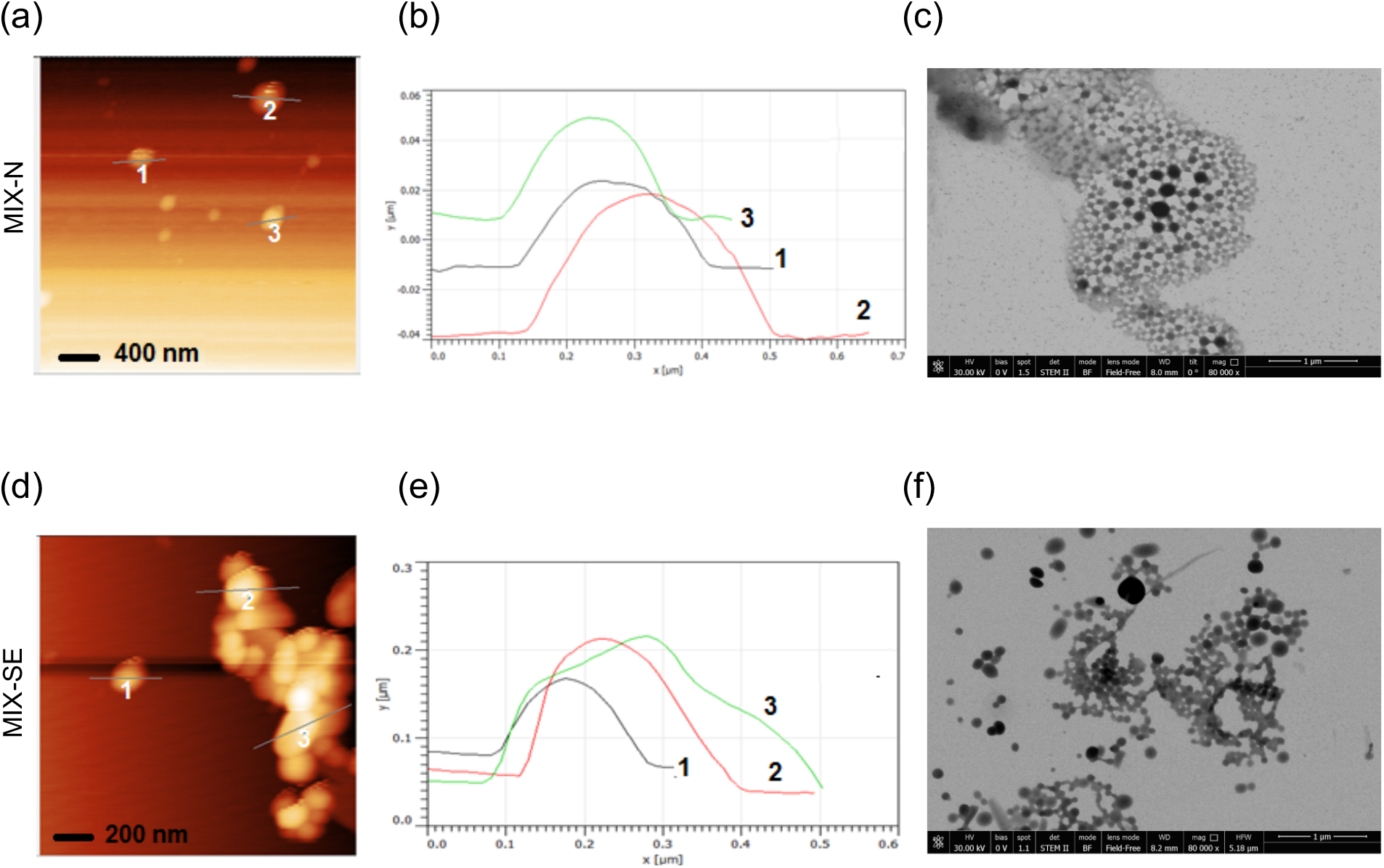
Microscopic characterization of hybrid-g7-NPs-chol. (a), (b) and (c) Atomic Force Microscopy (AFM) image (a) with relative analyses of profiles (b) and Scanning Transmission Electron microscopy (STEM) image (c) of MIX-N NPs. (d), (e) and (f) AFM image (d) with relative analyses of profiles (e) and STEM image (f) of MIX-SE NPs.

**Figure S2.**
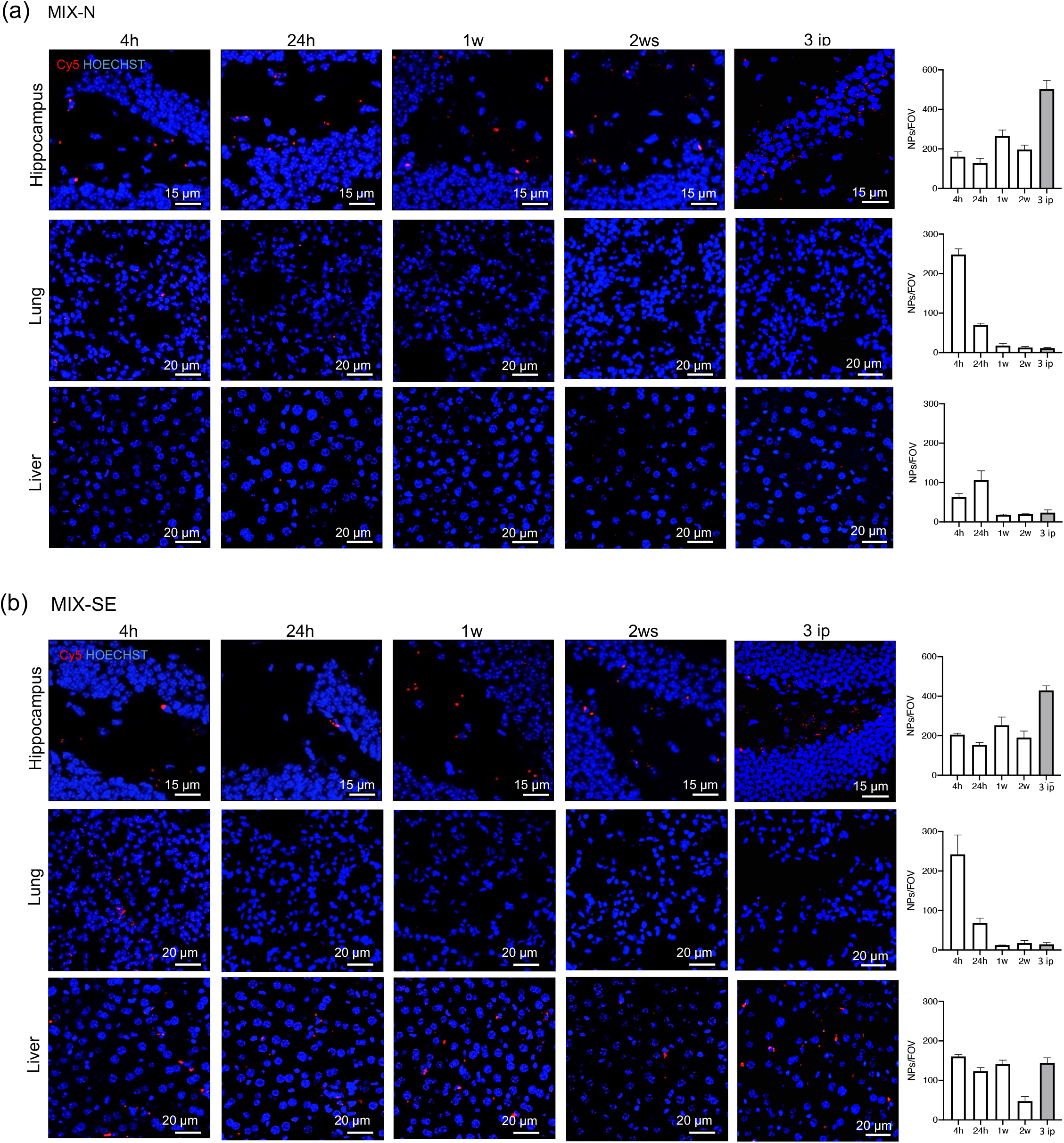
*In vivo* distribution of hybrid-cy5-g7-NPs-chol in other brain and peripheral tissues. (a) and (b). Representative confocal images of hippocampus, lung, and liver slices from wt mice that received 1 or 3 ip injection hybrid-Cy5-g7-NPs-chol (MIX-N in (a) or MIX-SE in (b)) and sacrificed after 4h, 24h, 1w, and 2w with relative quantification. Hoechst were used to counterstain nuclei (Ho, blue). Data are expressed as the number of g7-NPs-chol for 1 field of view ± standard error of the mean.

**Figure S3.**
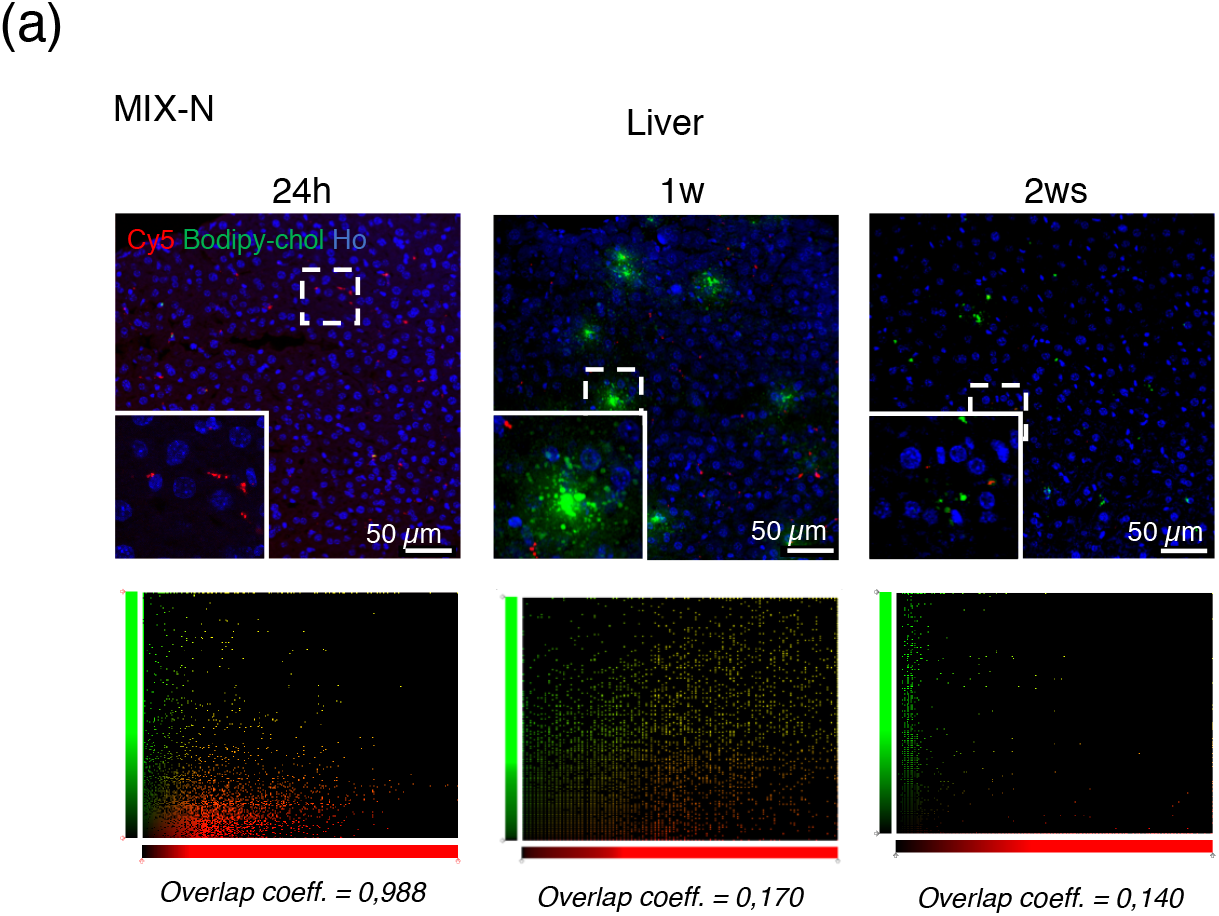
Cholesterol release from hybrid-Cy5-g7-NPs-bodipy-chol: qualitative analysis. Representative confocal image of liver brain slices from R6/2 mice ip injected with hybrid-Cy5-g7-NPs-bodipy-chol (MIX-N) and sacrificed after 24 h, 1 week or 2 weeks and relative co-localization of bodipy-chol and g7-NPs.

**Table S1.** Chemico-physical properties of NPs involved in the study and produced with different composition. Chemico-physical properties of NPs involved in the study and produced with different composition. In brackets SD. PDI stands for Polydispersity index, with values between 0 and 1. Lower is the PDI values, higher is the homogeneity of the sample in terms of size distribution. Amount of cholesterol is expressed in mg.

| Type | Sample Name |  | Size (nm) | PDI | Zeta Potential (mV) | Chol Content (mg Chol/100mg NPs) | Concentration (mg/mL) | In vivo experiments and analyses |
| --- | --- | --- | --- | --- | --- | --- | --- | --- |
| PLGA | PLGA-g7-NPs-chol |  | 180 (15) | 0.05 (0.02) | -22 (4) | 0.9 (0.1) | 9 | Mass spectrometry analysis |
| Hybrid | Cy5-g7-NPs-chol | MIX-N | 230 (12) | 0.16 (0.1) | -20 (5) | 28 (3) | 6.7 | Bio-distribution |
|  |  | MIX-SE | 250 (14) | 0.13 (0.5) | -20 (5) | 31 (3) | 7.4 | Bio-distribution |
|  | Cy5-g7-NPs-Bodipy-chol | MIX-N | 303 (4) | 0.25 (0.02) | -25 (4) | 32 (4) | 8.5 | qualitative studies of chol release |
|  |  | MIX-SE | 270 (15) | 0.16 (0.5) | -19 (6) | 26 (5) | 8.8 | qualitative studies of chol release |
|  | g7-NPs-d6-chol | MIX-N | 166 (4) | 0.22 (0.1) | -42 (1) | 24 (1) | 8.4 | d6-chol quantification by mass spectrometry |
|  | g7-NPs-chol | MIX-N | 240 (14) | 0.24 (0.04) | -25 (3) | 30 (5) | 8.6 | Mass spectrometry and behavioural analysis |

**Table S2.** Cholesterol release from hybrid-cy5-g7-NPs-bodipy-chol: overlap coefficient. Cholesterol release from hybrid-Cy5-g7-NPs-bodipy-chol: overlap coefficient.

| genotype | time after ip injection | overlap coefficient (MIX-N) |  |  | overlap coefficient (MIX-SE) |  |  |
| --- | --- | --- | --- | --- | --- | --- | --- |
|  |  | Cortex | Striatum | Liver | Cortex | Striatum | Liver |
| wt mice | 24h | 0.829 | 0.873 | 0.840 | 0.708 | 0.828 | N/D |
|  | 2ws | 0.667 | 0.544 | 0.055 | 0.542 | 0.647 | N/D |
| R6/2 mice | 24h | 0.738 | 0.799 | 0.988 | 0.888 | 0.873 | N/D |
|  | 2ws | 0.660 | 0.500 | 0.140 | 0.501 | 0.581 | N/D |

**Table S3.** Number of animals used in the study. Summary of all the trial performed and the animals used in this study.

| Trial number and date | Experimental groups | Bio-distribution | Chol release: quantitative analysis | Chol release: quantitative analysis | Mass spectrometry analysis | Behavioral analysis | Bioplex analysis |
| --- | --- | --- | --- | --- | --- | --- | --- |
|  |  | N mice | N mice | N mice | N mice | N mice | N mice |
| 1 - September 2017 | wt + hybrid-cy5-g7-NPs-chol MIX-N | 10 |  |  |  |  |  |
|  | wt + hybrid-cy5-g7-NPs-chol MIX-SE | 10 |  |  |  |  |  |
| 2 - November 2017 | wt + hybrid-cy5-g7-NPs-chol MIX-N | 10 |  |  |  |  |  |
|  | wt + hybrid-cy5-g7-NPs-chol MIX-SE | 10 |  |  |  |  |  |
| 3 - February 2018 | wt + hybrid-cy5-g7-NPs-bodipy-chol MIX-N |  | 9 |  |  |  |  |
|  | wt + hybrid-cy5-g7-NPs-bodipy-chol MIX-SE |  | 9 |  |  |  |  |
|  | R6/2 + hybrid-cy5-g7-NPs-bodipy-chol MIX-N |  | 9 |  |  |  |  |
|  | R6/2 + hybrid-cy5-g7-NPs-bodipy-chol MIX-SE |  | 9 |  |  |  |  |
| 4 - May 2018 | R6/2 + hybrid-g7-NPs-d6-chol MIX-N |  |  | 21 |  |  |  |
| 5 - January 2019 | wt saline |  |  |  | 3 | 10 |  |
|  | R6/2 saline |  |  |  | 3 | 8 |  |
|  | R6/2 + hybrid-g7-NPs-chol |  |  |  | 3 | 10 |  |
| 6 - November 2019 | wt saline |  |  |  |  | 5 |  |
|  | R6/2 saline |  |  |  |  | 4 | 4 |
|  | R6/2 + PLGA-g7-NPs-chol |  |  |  |  | 10 |  |
|  | R6/2 + hybrid-g7-NPs-chol |  |  |  |  | 7 | 5 |

